# Increased PHGDH expression uncouples hair follicle cycle progression and promotes inappropriate melanin accumulation

**DOI:** 10.1101/249250

**Authors:** Katherine R. Mattaini, Mark R. Sullivan, Allison N. Lau, Brian P. Fiske, Roderick T. Bronson, Matthew G. Vander Heiden

**Affiliations:** Koch Institute for Integrative Cancer Research, Cambridge, Massachusetts 02139, USA; Department of Biology, Massachusetts Institute of Technology, Cambridge, Massachusetts 02139, USA; Rodent Histopathology Core, Harvard Medical School, Boston, Massachusetts 02111, USA; Dana-Farber Cancer Institute, Boston, Massachusetts 02215, USA; Broad Institute, Cambridge, Massachusetts 02139, USA

**Keywords:** PHGDH, serine, melanin, hair follicle cycle, melanocyte

## Abstract

Copy number gain of the *PHGDH* gene that encodes the first enzyme of the serine biosynthesis pathway is found in some human cancers, including a subset of melanomas. In order to study the effect of increased *PHGDH* expression in tissues *in vivo*, we generated mice harboring a *PHGDH*^*tetO*^ allele that allows tissue-specific, doxycycline-inducible PHGDH expression. Tissues and cells derived from *PHGDH*^*tetO*^ mice exhibit increased serine biosynthesis. Histological examination of skin tissue from *PHGDH*^*tetO*^ mice reveals the presence of melanin granules in anagen II hair follicles, despite the fact that melanin synthesis is normally closely coupled to the hair follicle cycle and does not begin until later in the cycle. This phenotype occurs in the absence of any global change in hair follicle cycle timing. The inappropriate presence of melanin early in the hair follicle cycle following PHGDH expression is also accompanied by increased melanocyte abundance in anagen II skin. Together, these data support a model in which PHGDH expression affects melanocyte proliferation and/or differentiation and may provide insight into how PHGDH expression impacts normal melanocyte biology to promote melanoma.

**SIGNIFICANCE:** The significance behind copy number gain of *PHGDH* in human cancers is unclear. In this study, we generate a mouse model that mimics *PHGDH* gene copy number gain and characterize its effect on normal tissues. Increased PHGDH expression yields a phenotype of aberrant melanin production, which indicates that PHGDH expression may play a role in normal melanocyte biology. This result may provide insight into why *PHGDH* copy number gain is observed in melanoma more frequently than in most other tumor types.

## INTRODUCTION

D-3-phosphoglycerate dehydrogenase (PHGDH) is the first enzyme in the *de novo* serine biosynthesis pathway. Flux through this pathway can be important for the proliferation of some cancer cells, and the *PHGDH* gene is located in a region of focal genomic copy number gain that is associated with subsets of breast cancer and melanoma as well as cell lines derived from other cancer types (Locasale et al., 2011; Possemato et al., 2011). *PHGDH*-amplified cells are dependent on expression of catalytically active enzyme to proliferate (Mattaini et al., 2015). Furthermore, MCF-10A mammary epithelial cells with PHGDH overexpression show evasion of matrix detachment-induced cell death (anoikis), and high PHGDH expression is associated with negative clinical outcomes in breast cancer (Locasale et al., 2011; Pollari et al., 2011; Possemato et al., 2011), glioma (Liu et al., 2013) and cervical cancer (Jing, Heng, Aiping, Yafei, & Shulan, 2013). In addition to gene amplification, PHGDH expression can be upregulated through transcriptional and epigenetic mechanisms (Adams, 2007; Ding et al., 2013; Nilsson et al., 2012). However, to date, whether increased PHGDH expression in tissues promotes cancer initiation or progression and what impact increased enzyme activity has on normal physiology has not been studied.

Because *PHGDH* gene copy number gain is observed with higher frequency in melanoma compared to other cancers (Locasale et al., 2011; Possemato et al., 2011), the effect of PHGDH expression on melanocyte biology is of particular interest. Melanocytes are the main pigment-producing cells in mammals. The two most common melanin pigments generated by melanocytes, brown-black eumelanin and red-yellow pheomelanin, are both heterogeneous polymers. Unlike the vast majority of biosynthesis pathways in the body, many of the reactions of melanogenesis are spontaneous; therefore, local conditions heavily influence melanin structure and might be influenced by changes in metabolism (Hearing, 2011). The monomeric units of melanin are ultimately derived from the amino acid tyrosine, which is in turn synthesized from phenylalanine. Eumelanin is composed of 5,6-dihydroxyindole (DHI) and 5,6-dihydroxyindole-2-carboxylic acid (DHICA) units, and eumelanin synthesis utilizes the enzymes tyrosinase, tyrosinase-related protein-1 (Tyrp1) and dopachrome tautomerase (Dct), also known as tyrosinase-related protein-2 (Tyrp2). Pheomelanin is composed of benzothiazine units, derived from the reaction of the tyrosine product dopaquinone (DQ) with cysteine, and pheomelanin synthesis requires only tyrosinase activity (Hearing, 2011; Simon, Peles, Wakamatsu, & Ito, 2009). Melanin is produced in specialized lysosome-related organelles known as melanosomes, which are transferred to keratinocytes to ultimately pigment the skin or hair. Melanosomes undergo a stepwise maturation process, during which a proteinaceous matrix is formed, melanin synthesis enzymes become active, and melanin is synthesized and deposited (Schiaffino, 2010). Once melanosome maturation is complete, the melanosome is actively transferred to keratinocytes (Marks & Seabra, 2001). How each of these steps in melanogenesis would be affected by alterations in serine biosynthesis is unclear.

In mice, cutaneous melanocytes in truncal skin are exclusively follicular. Melanogenesis in follicular melanocytes is closely coupled to hair follicle (HF) cycling. Once a HF and the first hair are formed during morphogenesis, the entire base of the HF, the cycling portion, undergoes programmed cell death during a period known as catagen. The HF then enters a “resting” phase, telogen, before the anagen period (Chase, 1954; Fuchs, 2007) during which the entire lower portion of the HF is repopulated from epithelial and melanocyte stem cells located in the bulge region (Cotsarelis, Sun, & Lavker, 1990; Nishimura et al., 2002). Initiation of melanogenesis is tightly coupled to anagen progression (Slominski & Paus, 1993), with the first melanin granules visible in the HF during the anagen IIIa stage, when the hair follicle bulb extends to the border of the dermis and subcutis (Muller-Rover et al., 2001). Though serine biosynthesis is not obviously connected to HF cycling, serine biosynthesis pathway enzymes may affect differentiation of survival of stem cells (Hwang et al., 2016; Samanta et al., 2016), which could potentially perturb HF cycle progression.

To study how increased PHGDH expression affects normal tissue function in mice, we developed a transgenic mouse harboring a human PHGDH cDNA under the control of a doxycycline-inducible promoter. We found that expression of PHGDH results in premature appearance of melanin in HFs as well as an increased number of melanocytes in whole skin, suggesting that PHGDH expression affects melanocyte proliferation and/or differentiation, which may contribute to selection for increased PHGDH expression in cancer.

## RESULTS

### Generation of a *PHGDH*^*tetO*^ allele

In order to model the consequences of *PHGDH* amplification observed in cancer and study the effect of increased PHGDH expression in tissues, a transgenic mouse was engineered using a system designed by Beard, Hochedlinger, Plath, Wutz, and Jaenisch (2006). A human PHGDH cDNA under the control of the tetracycline operator minimal promoter (tetO) was integrated downstream of one copy of the mouse Col1A locus using flippase recombinase and flippase recognition targeting sites (FLP-FRT) in genetically modified KH2 ES cells (Supplemental Figure 1A). The KH2 ES cells also contain an *M2rtTA* (reverse tetracycline transactivator) allele under the control of the endogenous Rosa26 promoter that is active in most tissues. The presence of this *Rosa26-M2rtTA* allele allows PHGDH expression to be achieved in most cells and tissues upon doxycycline (dox) administration.

Targeted KH2 cell clones were screened by Southern blot for integration of the transgene into the ColA1 locus (Supplemental Figure 1B). Two clones contained one wild-type and one successfully targeted Col1A locus, and these were used to generate chimeric mice that were bred to acquire mice with germline transmission of the *PHGDH*^*tetO*^ allele. Southern blot analysis confirmed expected targeting of *PHGDH*^*tetO*^ into the ColA1 locus in these animals (Supplemental Figure 1C). To facilitate animal husbandry and analysis, we refined a widely-utilized PCR-based genotyping strategy to determine *PHGDH*^*tetO*^ and *Rosa26-M2rtTA* genotypes in these mice (Supplemental Figure 1D)(The Jackson Laboratory, 2016).

### Characterization of *PHGDH*^*tetO*^ mice

PHGDH is only expressed in tissues from mice with both the *PHGDH*^*tetO*^ and the *Rosa26-M2rtTA* alleles and only upon exposure of tissues to doxycycline (dox) (Figure 1A). To test whether increased PHGDH expression affects viability, breeding pairs of *PHGDH*^*tetO*^ hemizygotes were continually fed a diet containing dox to induce PHGDH expression in the whole body. Offspring from these crosses were born in expected Mendelian ratios (Supplemental Figure 1E). Expression from the Rosa26 promoter is active by the blastocyst stage of the developing embryo (Zambrowicz et al., 1997), and dox readily crosses the placenta to regulate transgene expression in the developing embryo (Fedorov, Tyrsin, Krenn, Chernigovskaya, & Rapp, 2001; Perl, Tichelaar, & Whitsett, 2002; Perl, Wert, Nagy, Lobe, & Whitsett, 2002; Shin, Levorse, Ingram, & Tilghman, 1999), so this result suggests that increased PHGDH expression in the whole embryo has no effect on viability.

**Figure 1.**
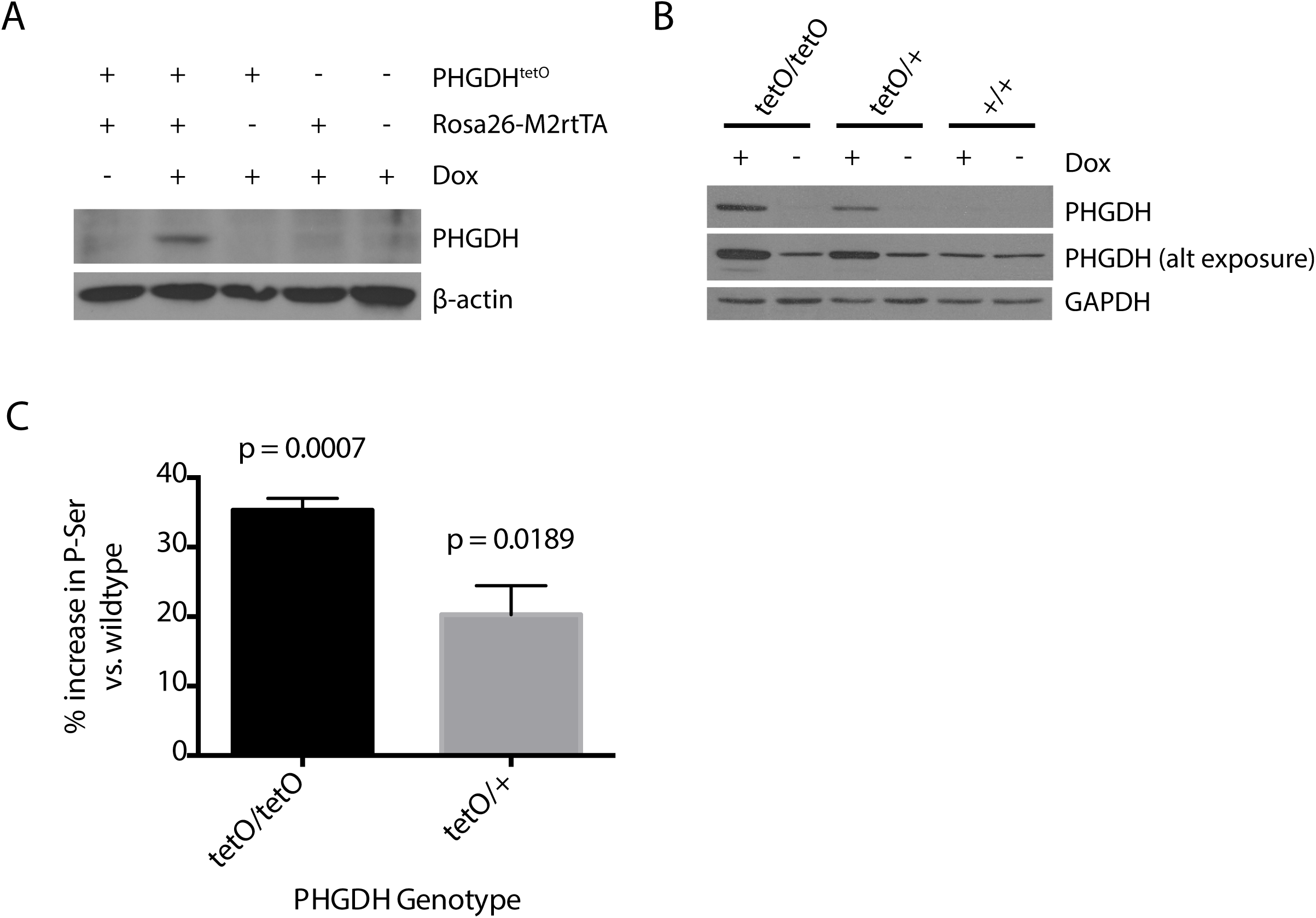
Introducing a *PHGDH*^*tetO*^ allele into mice increases PHGDH expression and serine synthesis. **(A)** Western blot analysis to assess PHGDH expression in liver lysates from mice exposed to a doxycycline containing diet (Dox) or a control diet for 5 days. β-actin expression was also assessed as a loading control. (B) Western blot analysis to assess PHGDH expression in MEFs derived from *PHGDH*^*tetO*^ mice that were cultured in media with or without doxycycline (Dox) as for 72 hours as indicated. Both a light and dark exposure (alt exposure) is shown, as is GAPDH expression as a loading control. (C) The percent increase in the concentration of intracellular phosphoserine (P-ser) in MEFs derived from *PHGDH*^*tetO*^ mice harboring one (tetO/+) or two (tetO/tetO) transgene alleles relative to levels found in MEFs derived from wildtype mice is shown. All MEFs were cultured for 4 days in media with doxycycline prior to measurement of P-Ser levels by LC-MS. Data shown represent the mean (+ SEM. The increase is statistically significant with p values from two-tailed Student’s T test.

Embryonic fibroblasts (MEFs) derived from *PHGDH*^*tetO*^*;Rosa26-M2rtTA* mice (referred to hereafter as *PHGDH*^*tetO*^ mice) display dose-dependent, dox-inducible PHGDH expression (Figure 1B). The antibody used throughout this study recognizes both human and mouse PHGDH proteins with roughly equal affinity by Western blot (Supplemental Figure 1F); thus, PHGDH expression observed in *PHGDH*^+/+^ MEFs and in conditions without dox-induced transgene activation reflect mouse PHGDH protein expressed from the endogenous locus. In previous studies examining a variety of cell lines and tissues, expression of PHGDH at the protein level correlates with serine biosynthesis pathway flux (Davis & Fallon, 1970; Locasale et al., 2011; Possemato et al., 2011). Similar results are obtained following transgene expression, as dox-treated *PHGDH*^*tetO*^ MEFs show a dose-dependent increase in both PHGDH protein and the concentration of the unique serine biosynthesis pathway intermediate phosphoserine compared to dox-treated wildtype MEFs (Figure 1 B-C). These data suggest the transgene expression can increase serine biosynthesis in cells.

### Mice with long-term PHGDH overexpression are grossly normal

*PHGDH*^*tetO*^ mice were exposed to dox diet beginning at 6 weeks of age and maintained on this diet for 16-18 months. During this time, mice were monitored weekly without evidence of any obvious abnormalities while alive and at necropsy. Liver and skin samples were analyzed by Western blot for PHGDH protein expression. Some samples showed less PHGDH expression than expected after 16-18 months of dox exposure (Supplemental Figure 2), but liver and skin samples from the same individual showed consistent expression levels suggesting that differences in transgene silencing might underlie the variability in expression. Histological analysis of skin, brain, white and brown fat, mammary gland, pancreas, liver, spleen, kidney, colon, lung and heart tissue in this cohort from control mice and *PHGDH*^*tetO*^ mice with high PHGDH expression by Western blot was unremarkable; however, analysis of skin in younger animals suggested an abnormality was present in this tissue.

### Pre-anagen IIIa hair follicles in *PHGDH*^*tetO*^ mice inappropriately contain melanin granules

Because *PHGDH* amplification is associated with some melanomas (Locasale et al., 2011), melanocytes were a cell type of interest when monitoring for phenotypes in the *PHGDH*^*tetO*^ mice. When examining the skin of 3.5 month-old mice treated with dox for 9 days, an anomaly in follicular melanin was observed (Figure 2A). During the normal hair follicle (HF) cycle, melanin is not visible in the hair bulb until anagen IIIa, when the bulbs are characteristically located within the border between the dermis and the subcutis (Muller-Rover et al., 2001). The bulbs of the HFs pictured in Figure 2A are surrounded entirely by dermis, identifying them as pre-anagen IIIa; however, in *PHGDH*^*tetO*^ HFs melanin granules are clearly visible. Ordinarily, during catagen, all cells from the cycling portion of the HF undergo apoptosis, including the melanocytes. Any melanin they have produced is passed to the keratinocytes that make up the hair itself, so that melanin is no longer present in the bulb before new melanin is produced in anagen IIIa of the next HF cycle. Occasionally, melanin granules produced in a previous HF cycle will not be extruded with the hair shaft and are visible in the dermal papilla in telogen, anagen I, or anagen II (Tobin, Hagen, Botchkarev, & Paus, 1998). However, the *PHGDH*^*tetO*^ skin had a significantly greater proportion of pre-anagen IIIa HFs displaying melanin than the wildtype skin (Figure 2B). Furthermore, although some pre-anagen IIIa HFs in the wildtype skin displayed one or two melanin granules, almost none had three or more (Figure 2C). In contrast, many melanin-containing follicles in the *PHGDH*^*tetO*^ skin had as many as 5-10 granules.

**Figure 2.**
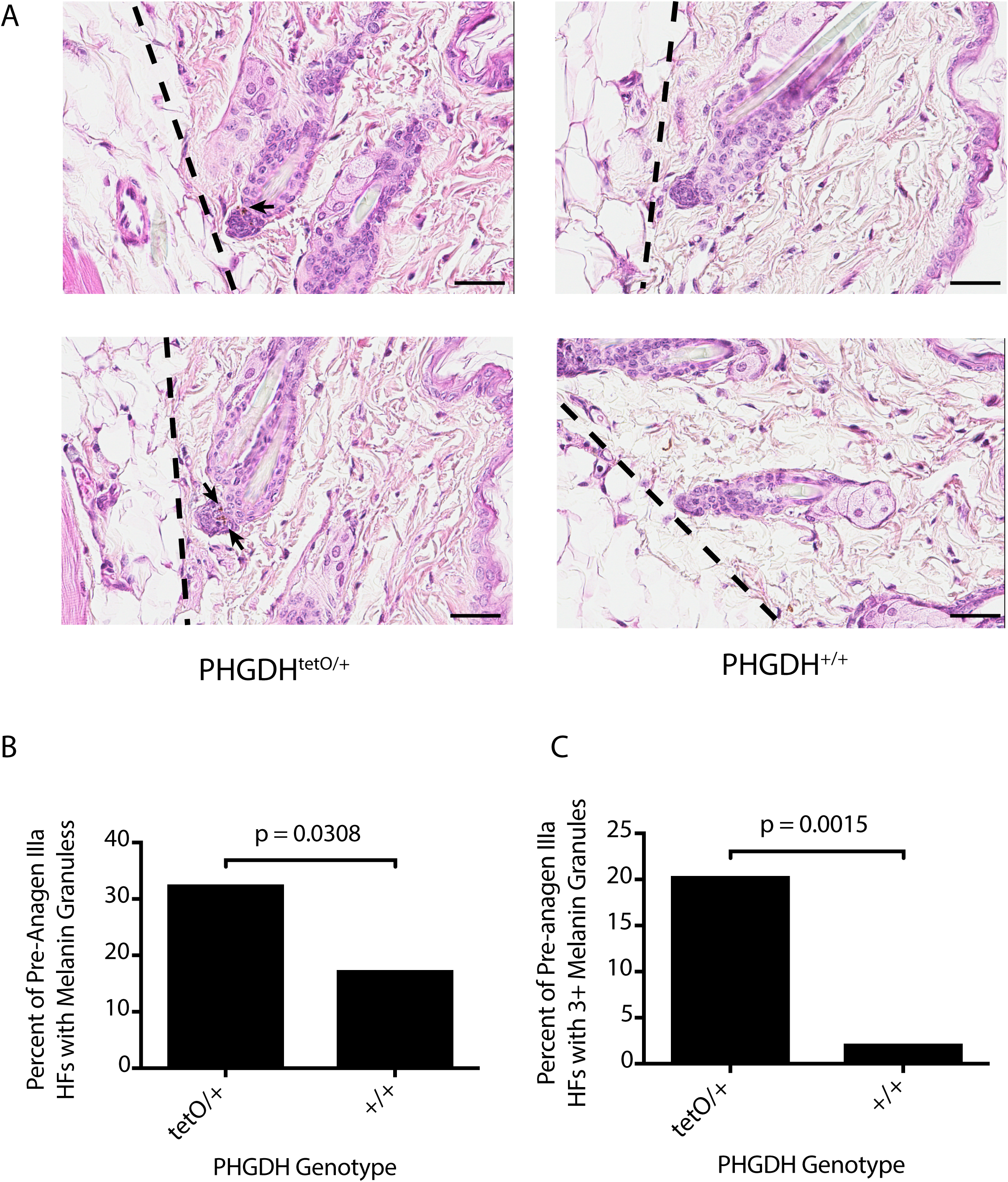
Pre-anagen IIIa hair follicles in *PHGDH*^*tetO*^ mice contain melanin granules. **(A)** Representative H&E staining of skin sections from 3.5-month-old mice of the indicated genotypes that had been exposed to a doxycycline-containing diet for 9 days. Dotted lines delineate the border between the dermis and the subcutis. Arrow indicates melanin granules in the hair follicles (HFs) of the *PHGDH*^*tetO*^ mouse. All hair follicle shown are pre-anagen IIIa as they are contained completely within the dermis. Images were obtained at 40x magnification. Scale bar = 30 μm. **(B)** Quantitation of the percent of pre-anagen IIIa hair follicles (HF) in each genotype that contain any melanin granules. Data shown represent the % observed when analyzing 167 HFs from one PHGDH^tetO^ mouse and 46 HFs from one wild type mouse. **(C)** Quantitation of the percent of pre-anagen IIIa hair follicles (HF) in each genotype with three or more melanin granules. Data shown represent the % observed when analyzing 167 HFs from one PHGDH^tetO^ mouse and 46 HFs from one wild type mouse. The percent increase in hair follicles with melanin granules shown in (B) and (C) is statistically significant with p values derived from one-tailed Fisher’s exact test.

### PHGDH expression does not globally affect timing of the hair follicle cycle

To further characterize this phenotype, HF cycling was synchronized by plucking hairs from a region of skin to induce HFs in that region to enter a new cycle. Skin was then harvested at defined time points to examine a desired cycle stage (Muller-Rover et al., 2001). In order to determine the effect of PHGDH overexpression on follicular melanin throughout the HF cycle, two approaches were used: one set of mice was fed dox diet for two days before plucking (red bar) and the other set for thirty days before plucking (blue bar) to synchronize the HF cycle (Figure 3A). The first few HF cycles following birth are relatively synchronous across individuals (Muller-Rover et al., 2001). Therefore, a 30-day pre-induction with dox following by plucking at 49 days of age allows PHGDH overexpression during the entire cycle preceding plucking, from telogen to telogen. Conversely, the 2-day pre-induction only allowed PHGDH overexpression during the very end of the HF cycle preceding synchronization. By using two different pre-induction times, we aimed to determine whether the melanin phenotype required PHGDH overexpression in only the current HF cycle or if expression in the preceding cycle was required for melanin accumulation.

**Figure 3.**
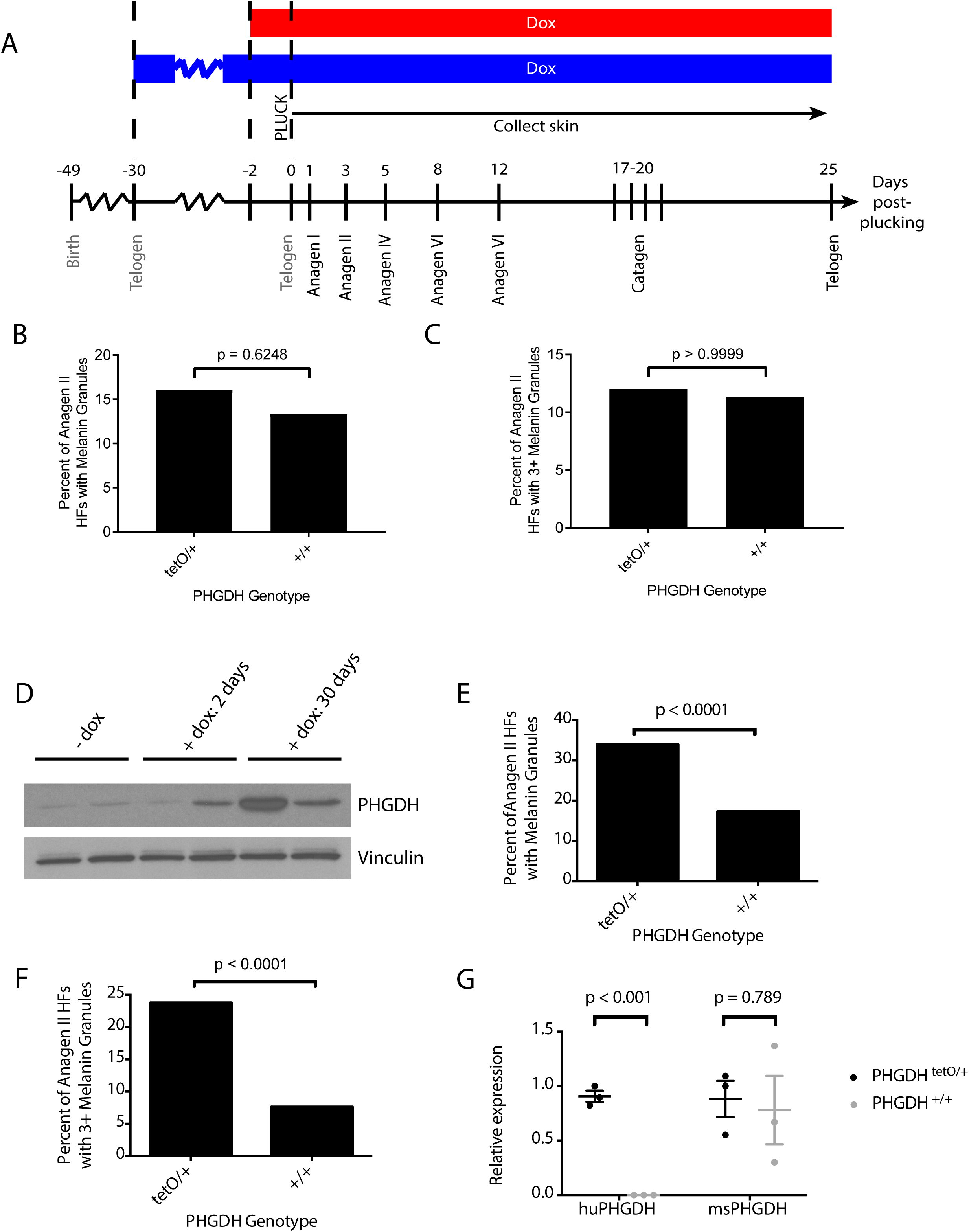
PHGDH expression during the previous hair follicle cycle leads to increased melanin accumulation. **(A)** A region of hair was plucked from 49 day-old mice (at the second telogen) to synchronize the hair follicle cycle, and skin samples were collected at defined days thereafter. Data were collected from *PHGDH*^*tetO*^ or control (+/+) mice that were exposed to doxycycline (Dox) for either 2 days or 30 days prior to synchronization. Shown is a schematic of the experiment, with the red bar depicting mice exposed to doxycycline diet for 2-days before synchronization, and the blue bar depicting mice exposed to doxycycline for 30-days before synchronization. **(B)** Quantitation of the percent of anagen II hair follicles (HFs) containing any melanin granules in mice of each genotype exposed to doxycycline for 2 days prior to synchronization. Data shown represent the % observed when analyzing 50 HFs per mouse from 3 mice of each genotype **(C)** Quantitation of the percent of anagen II hair follicles with three or more melanin granules in mice of each genotype exposed to doxycycline for 2 days prior to synchronization. Data shown represent the % observed when analyzing 50 HFs per mouse from 3 mice of each genotype No statistically significant increase in hair follicles with melanin granules were observed in (B) or (C) with p-values derived from two-tailed Fisher’s exact test. **(D)** Western blot analysis for PHGDH expression in skin from mice never exposed to doxycycline-containing diet (-dox) or fed a doxycycline-containing diet for 2 or 30 days as indicated. Vinculin expression is also shown as a loading control. **(E)** Quantitation of the percent of anagen II hair follicles (HFs) containing any melanin granules in mice of each genotype exposed to doxycycline for 30 days prior to synchronization. Data shown represent the % observed when analyzing 50 HFs per mouse from 3 mice of each genotype. **(F)** Quantitation of the percent of anagen II hair follicles with three or more melanin granules in mice of each genotype exposed to doxycycline for 30-days prior to synchronization. Data shown represent the % observed when analyzing 50 HFs per mouse from 3 mice of each genotype. The increase in hair follicles with melanin granules shown in (E) and (F) is statistically significant with p-values derived from two-tailed Fisher’s exact test. **(G)** qPCR to assess species-specific PHGDH expression in anagen II skin isolated from mice of the indicated genotype exposed to doxycycline for 30-days prior to synchronization. An increase in human PHGDH (huPHGDH), but not mouse PHGDH (msPHGDH) expression is statistically significant with p values derived from unpaired Student’s t test. Data shown represent the mean (+/-SD).

Examination of skin at various time points after HF synchronization in mice exposed to dox for thirty days prior to plucking suggested PHGDH overexpression does not globally affect timing of the HF cycle (Supplemental Figure 3A). HFs in synchronized skin from both control and *PHGDH*^*tetO*^ mice were found in the expected stages for their collection days. Additionally, no perceptible differences visible by H&E staining were evident in any HF stage other than anagen II. The fact that the phenotype is anagen II-specific likely explains why it was not detected in the cohort of aged mice. Anagen II is relatively short compared to the entire HF cycle; thus, anagen II HFs are not abundant in mice of any age. Furthermore, HF cycling becomes more asynchronous as mice age (Paus & Foitzik, 2004), so that the likelihood of collecting a skin sample by chance with an abundance of anagen II HFs is further decreased.

### Anagen II hair follicles in synchronized *PHGDH*^*tetO*^ skin contain melanin granules

Anagen II follicles in synchronized skin of *PHGDH*^*tetO*^ mice with only a 2-day dox pre-induction show the presence of inappropriate melanin granules, but the neither the proportion of HFs with melanin (Figure 3B) nor the fraction of HFs with three or more melanin granules (Figure 3C) is significantly different than in wild-type mice. Analyzing skin by Western blot shows that 2-day pre-induction is sufficient to moderately increase PHGDH levels in some mice; however, the change in expression is higher in the skin of mice exposed to dox diet for 30 days (Figure 3D), raising the possibility that the absence of a melanin phenotype after 2 days of dox pre-induction is due to the latency of PHGDH expression.

Examination of anagen II skin from mice with a 30-day pre-induction showed inappropriate melanin accumulation to a degree that reproduced the initial phenotype observed (Supplemental Figure 3B). The anagen II HFs from the skin of *PHGDH*^*tetO*^ mice contained melanin more frequently (Figure 3E) and were more likely to have a high number of melanin granules (Figure 3F) than their wild-type counterparts. The percentages observed in this experiment were similar to those observed in the initial unsynchronized experiment (Figure 2B-C). To confirm that melanin accumulation is associated with increased PHGDH expression from the transgene, we designed qPCR primers specific for human or mouse PHGDH cDNA (Supplemental Figure 4A-D) and found that in synchronized *PHGDH*^*tetO*^ skin, expression of human PHGDH was increased while expression of mouse PHGDH was unchanged (Figure 3G).

### Melanin accumulation in *PHGDH*^*tetO*^ mice is caused by cell autonomous PHGDH expression and is not dependent on PHGDH overexpression in catagen

In order to determine whether the melanin phenotype observed in the *PHGDH*^*tetO*^ mice is cell autonomous, we crossed *PHGDH*^*tetO*^ mice to mice harboring a *Dct-rtTA* allele that allows melanocyte-specific transgene expression (Zaidi, Davis, et al., 2011; Zaidi, Hornyak, & Merlino, 2011). With a 30-day pre-induction, skin from *PHGDH*^*tetO*^*;Dct-rtTA* mice displayed melanin granules in anagen II HFs with similar percentages as those observed in mice with a *Rosa26-M2rtTA* allele (Figure 4A-B), suggesting that the PHGDH-dependent presence of melanin in anagen II HFs is a melanocyte-autonomous event.

**Figure 4.**
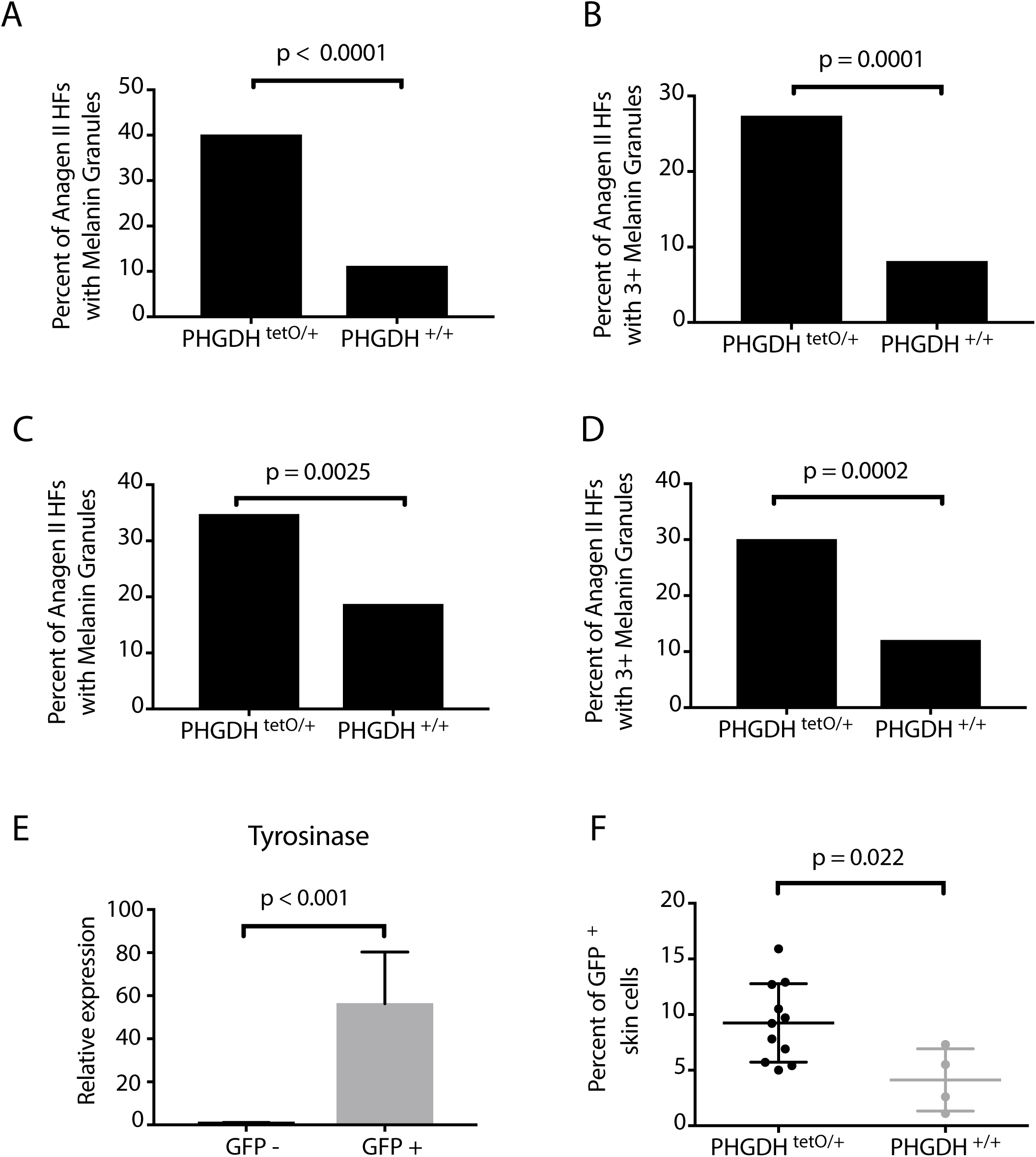
Increased PHGDH expression in melanocytes drives melanin accumulation in anagen II hair follicles and increases melanocyte abundance. **(A)** *PHGDH*^*tetO*^ mice were crossed to *Dct-rtTA* mice to drive increased PHGDH expression solely in melanocytes. Quantitation of the percent of anagen II hair follicles (HFs) containing any melanin granules in skin from mice of the indicated genotype exposed to doxycycline for 30 days prior to hair follicle synchronization as described in Figure 3. Data shown represent the % observed when analyzing 50 HFs per mouse from 3 mice of each genotype. **(B)** Quantitation of the percent of anagen II hair follicles (HFs) with three or more melanin granules in in skin from mice of the indicated genotype exposed to doxycycline for 30 days prior to hair follicle synchronization. Data shown represent the % observed when analyzing 50 HFs per mouse from 3 mice of each genotype. **(C)** Quantitation of the percent of anagen II hair follicles (HFs) containing any melanin granules in skin from mice of the indicated genotype exposed to doxycycline for 2 days prior to hair follicle synchronization. Data shown represent the % observed when analyzing 50 HFs per mouse from 3 mice of each genotype **(D)** Quantitation of the percent of anagen II hair follicles (HFs) with three or more melanin granules in in skin from mice of the indicated genotype exposed to doxycycline for 2 days prior to hair follicle synchronization. Data shown represent the % observed when analyzing 50 HFs per mouse from 3 mice of each genotype. The increase in hair follicles with melanin granules shown in (A-D) is statistically significant with p-values derived from twotailed Fisher’s exact test. **(E)** qPCR to assess tyrosinase expression (a melanocyte-specific enzyme) using of isolated from GFP‐ and GFP+ cells isolated in **(F)**. Data shown represent the mean (+/-SD). The increase in tyrosinase expression is significant with p-values derived from an unpaired Student’s t test. **(F)** *PHGDH*^*tetO*^; *Dct-rtTA* mice were crossed to *H2B-GFP*^*tetO*^ mice such that melanocytes would express both PHGDH and GFP. After exposure of these mice and control mice to doxycycline for 30-days prior to hair follicle synchronization, anagen II skin samples were isolated from animals of the indicated genotype and subjected to FACS analysis to assess melanocyte abundance in the skin. Data shown represent the mean (+/-SD). The increase in GFP+ melanocytes in hair follicles from PHGDH^tetO/+^ mice is statistically significant with p-values derived from an unpaired Student’s t test.

To evaluate whether PHGDH expression is required in the previous HF cycle for this phenotype, we used a 2-day pre-induction with dox. Though a 2-day pre-induction led to only weak PHGDH expression when driven by *Rosa26-M2rtTA*, the melanocyte specific *Dct-rtTA* is predicted to promote higher PHGDH expression in these cells. Indeed, we found that with a 2 day pre-induction, skin from *PHGDH*^*tetO*^;*Dct-rtTA* mice displayed melanin granules in anagen II HFs at a higher rate than skin from wildtype mice (Figure 4 C-D). The presence of the melanin phenotype with a 2-day pre-induction suggests that the phenotype does not depend on PHGDH overexpression during the previous catagen.

This argues against PHGDH promoting survival of melanocytes that would normally die during the previous catagen phase. Instead, the effect of PHGDH expression on uncoupling melanin appearance with the normal HF cycle progression only requires the presence of PHGDH during the earliest phases of the HF cycle.

### Increased PHGDH expression in melanocytes increases melanocyte abundance in anagen II skin

To determine whether the presence of excess melanin granules in anagen II HFs is related to a change in melanocyte number, we designed an approach to quantify melanocyte abundance using flow cytometry. *PHGDH*^*tetO*^; *Dct-rtTA* mice were crossed to *H2B-GFP*^*tetO*^ mice (Tumbar et al., 2004; Zaidi, Davis, et al., 2011; Zaidi, Hornyak, et al., 2011) so that melanocytes would express both PHGDH and GFP. The resulting mice were then exposed to a dox diet for 30 days, plucked, and skin was collected in anagen II. Adapting previously described protocols (Joshi et al., 2017; Zaidi, Davis, et al., 2011), this skin was then dissociated into single-cell suspension and sorted by flow cytometry into GFP-positive and GFP-negative populations in order to quantitate the effect of PHGDH expression on the relative abundance of GFP-positive melanocytes (Supplemental Figure 5A). In order to validate that the GFP-positive cells were indeed melanocytes, we performed qPCR for tyrosinase, a melanocyte marker, which was present in GFP-positive cells, and nearly undetectable in GFP-negative cells (Figure 4E). Conversely, expression of KPRP, a keratinocyte marker, and AdipoQ, an adipocyte marker, were restricted to the GFP-negative cells (Supplemental Figure 5B-C). The proportion of GFP-positive cells was significantly higher in anagen II skin from mice with the *PHGDH*^*tetO*^ allele (Figure 4F), suggesting that melanocytes are more abundant in anagen II skin when PHGDH is overexpressed.

## DISCUSSION

High *PHGDH* expression is observed in select cancer cells and in some cases is necessary for proliferation and survival (Locasale et al., 2011; Possemato et al., 2011). Genomic copy number gain involving *PHGDH* is observed with higher frequency in melanoma than in other cancers (Locasale et al., 2011), and in this regard it is interesting that increased PHGDH expression driven by a ubiquitous promoter in mice results in a phenotype involving melanocytes such that progression of the hair follicle cycle is uncoupled from melanin appearance. This effect on normal melanocyte biology may provide insight into how PHGDH expression contributes to melanoma.

A key unanswered question is why hair follicles in *PHGDH*^*tetO*^ mice display an increased number of melanin granules and melanocytes early in the HF cycle. Unlike human epidermal melanocytes, which perform melanogenesis constantly, melanogenesis in follicular pigment cells is invariably linked to the HF cycle and no genetic modifications in mice have been reported that induce new melanin synthesis in a catagen or telogen follicle (Paus & Foitzik, 2004). There are a multitude of ways in which PHGDH expression might affect melanin appearance. Recent mathematical modeling of cycles of varying lengths in individual hair follicles has suggested that HF exist in two quasi-steady states, growth/anagen and rest/telogen, and that the transition between the two stages is stochastic and based on a threshold effect of integrated signals (Al-Nuaimi, Goodfellow, Paus, & Baier, 2012; Bernard, 2012; Halloy, Bernard, Loussouarn, & Goldbeter, 2002). The signaling pathways that are integrated to determine this threshold are numerous, and include the Wnt/β-catenin, BMP/TGFβ, Shh, Notch, EGF/FGF (Lee & Tumbar, 2012) and mTOR pathways (Deng et al., 2015). PHGDH has no known connections with these pathways; however all have been implicated in cancer.

Melanogenesis itself involves the production, survival and differentiation of melanocytes (Tobin, 2008), functional melanosome biogenesis (Schiaffino, 2010), appropriate transcription, translation, modification, and activity of synthetic enzymes such as tyrosinase (Videira, Moura, & Magina, 2013; Wang & Hebert, 2006), input from autocrine and paracrine signals (Hsiao & Fisher, 2014), and availability of substrate and appropriate chemical conditions for melanogenesis, including pH and redox state (Hearing, 2011; Schallreuter, Kothari, Chavan, & Spencer, 2008). Increased serine synthesis might affect one or more of these processes in melanocytes or melanocyte stem cells. Alterations in redox state may be relevant to PHGDH expression, as increased serine synthesis is associated with resistance to oxidative stress in melanoma and breast cancer (Piskounova et al., 2015; Samanta et al., 2016). PHGDH could also affect follicular melanogenesis through an indirect effect, such as promotion of inappropriate differentiation of melanocytic stem cells into melanocytes or proliferation of those melanocytes.

Regardless of the effects on specific signaling pathways, an important question is whether melanocytes in the *PHGDH*^*tetO*^ mice display altered proliferation, differentiation or cell death, as this may be relevant to understanding why PHGDH is frequently amplified in melanoma. We found that driving expression of PHGDH exclusively in melanocytes via the *Dct-rtTA* allele led to an increased abundance of melanocytes in anagen II skin at the same time that we observe the melanin phenotype. There is precedent for PHGDH overexpression allowing evasion of cell death, as MCF-10A cells in the center of a 3D acinar structure inappropriately survive with PHGDH overexpression (Locasale et al., 2011). Differentiated, pigment-producing melanocytes normally undergo apoptosis during catagen (Tobin et al., 1998). Apoptosis of melanogenic pigment cells during catagen is thought to protect against transformation, since melanogenesis is highly oxidative, and continual melanin synthesis is proposed to cause DNA damage and promote cellular transformation (Tobin, 2008). However, we found that the melanin phenotype in the *PHGDH*^*tetO*^ mice did not require PHGDH expression during the previous catagen. Furthermore, telogen HFs following the 30-day pre-induction did not display significant melanin accumulation in either *PHGDH*^*tetO*^ or control mice, which one would expect to see if melanocytes were evading apoptosis. Together, these data suggest that the increase in melanocyte number due to PHGDH expression is likely related to increased proliferation and/or changes in differentiation.

Long-term expression of PHGDH based on ubiquitous Rosa26-rtTA expression revealed no gross phenotype save an increase in melanin granules in anagen II hair follicles. Importantly, the lack of tumor formation argues that PHGDH expression alone may not be sufficient to drive cancer. However, although it is relatively ubiquitous in expression, the *Rosa26-M2rtTA* allele does not drive expression to high levels in all tissues (Zambrowicz et al., 1997). It is possible that crossing the *PHGDH*^*tetO*^ mouse to mice with other tissue-specific rtTA alleles driving higher expression, or combining PHGDH expression with other genetic events could lead to additional phenotypes or promote cancer.

## METHODS

All mouse studies were performed in accordance with institutional guidelines and approved by the MIT Committee on Animal Care.

### Generation of *PHGDH*^*tetO*^ Mice

The system used to generate the *PHGDH*^*tetO*^ mice was developed and described by Beard et al. (2006). Human PHGDH cDNA with GenBank Accession BC011262.1 from Open Biosystems (MHS1010-73507) was amplified with the following primers:

PHGDH MfeI F: 5’-CAATTGGCCACCATGGCTTTTGCAAATCTGCGGAAAGT-3’

PHGDH Mfe R: 5’-CAATTGTTAGAAGTGGAACTGGAAGGCTTCAG-3’

This insert was digested with MfeI from NEB (R0589) and cloned into the EcoRI sites in the pgk-ATG-frt plasmid from Addgene (#20734) to generate a targeting plasmid using standard molecular biology techniques. Sequencing was used to screen for the correct insert orientation and confirm cDNA sequence. The targeting plasmid was co-electroporated with pCAGGS-flpE plasmid (Addgene, # 20733) into F1 C57BL/6 x 129S4 hybrid KH2 ES cells. The KH2 cells as well as pgk-ATG-frt and pCAGGS-flpE-puro were kind gifts from Rudolf Jaenisch (plasmids via Addgene). Clonal selection of ES cells was performed with 150 ug/ml hygromycin B for 9 days, and 8 individual clones were screened by Southern blot as described below. Two ES clones with a properly integrated PHGDH transgene in the Col1a1 locus were injected independently into C57BL/6 blastocysts to produce chimeric mice. The chimeric C57BL/6 x 129S4 *PHGDH*^*tetO*^ transgene founder mice were mated to C57BL/6 background and some showed germline transmission. Before this study and over the course of these experiments, the mice were continually backcrossed onto the C57BL/6 background.

### Southern Blotting

Genomic DNA was digested with SpeI from NEB (R0133). Digested DNA was then separated on an agarose gel, and neutral transfer was performed overnight using Hybond-XL membrane from GE Healthcare Biosciences (RPN303S). Membrane was crosslinked using a Stratalinker UV Crosslinker from Stratagene. The membrane was incubated with Stratagene QuickHyb Hybridization Solution from Agilent (201220). The probe was prepared from the Col1a-3'probe plasmid from Addgene (#20731) by digesting with XbaI and PstI from NEB (R0145 and R0140) and gel purifying the released probe. Purified probe was denatured, then labeled using α-^32^P-dCTP from PerkinElmer Life Sciences (BLU013H) and the Rediprime II DNA Labeling System from GE Healthcare Life Sciences (RPN1633) according to kit instructions. Labeled probe was then purified with Micro Bio-Spin P-6 Gel Columns from Bio-Rad (#732-6200) according to company instructions. Purified, labeled probe was mixed with salmon sperm DNA from Stratagene (201190). Immediately before using, probe was denatured. The probe was then incubated with the membrane and hybridization solution for 1 hr at 68°C. The membrane was washed, then exposed to autoradiography film in a cassette with an intensifier screen before developing.

### PCR Genotyping

PCR genotyping was performed using standard molecular biology techniques under conditions described below.

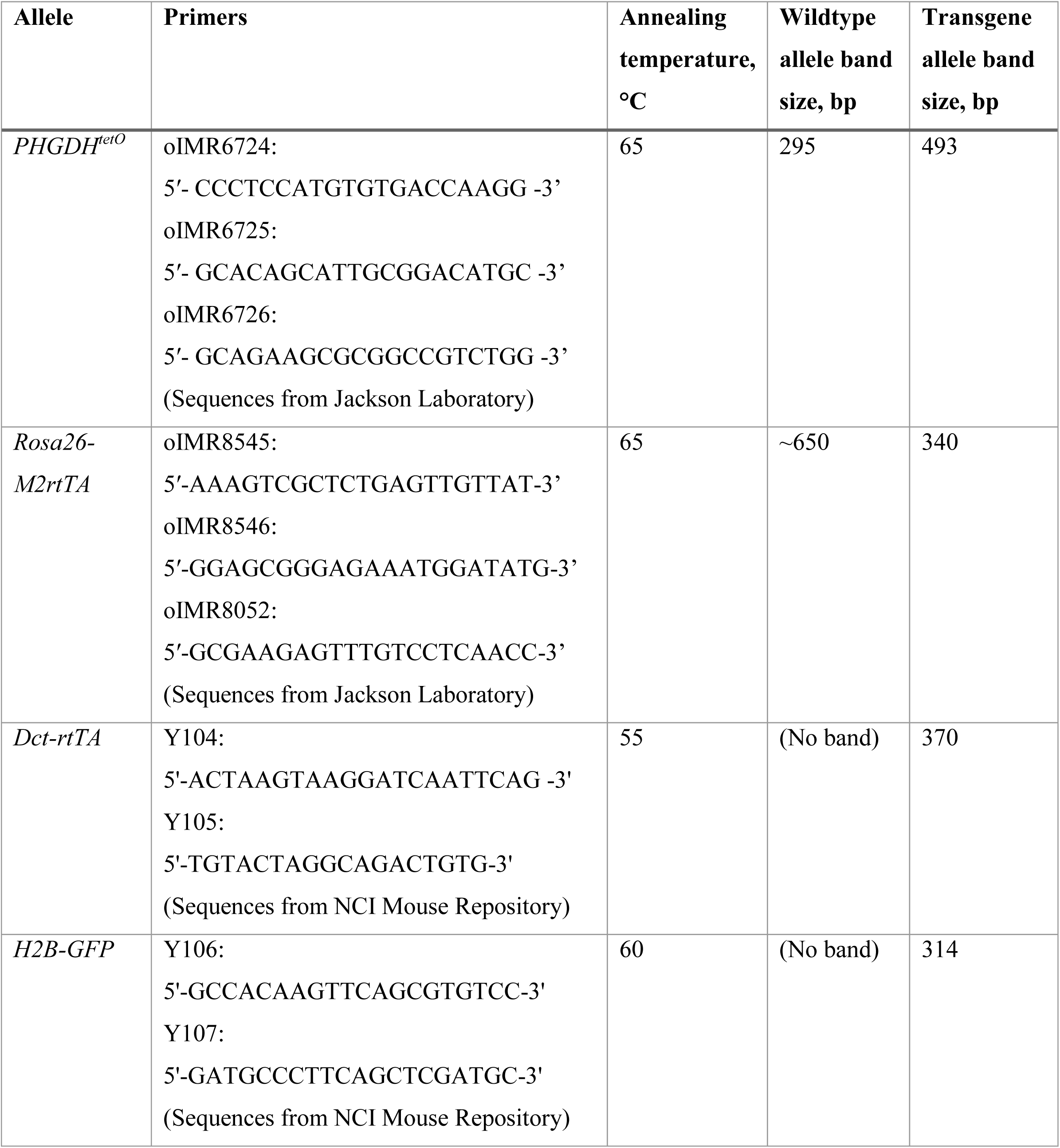

### Western Blotting

Western blots were performed using standard techniques with primary antibodies against PHGDH (Sigma, HPA021241), β-actin (abcam, ab1801), GAPDH (Cell Signaling Technology, 2118S), Hsp90 (Cell Signaling Technology, #4877), or vinculin (abcam, ab18058) and detected using HRP-conjugated secondary antibodies and chemiluminescence.

### Generation of Embryonic Fibroblasts & Cell Culture

MEFs were prepared from E13.5 *PHGDH^tetO/tetO^*, *PHGDH*^*tetO*/*+*^ or *PHGDH*^+/+^ embryos with the *Rosa26-M2rtTA* allele using standard protocols. MEFs were maintained in DMEM with pyruvate, 10% tet-free FBS, 2 mM glutamine, penicillin/streptomycin and 3.5 μl beta-mercaptoethanol per 500 ml DMEM.

### Mass Spectrometry Measurement of Phosphoserine

MEFs were grown in medium supplemented with 1 μg/ml doxycycline for 4 days before extraction. Cells were extracted in ice cold 1:4:5 water:methanol:chloroform with valine-D8 as an internal standard. The aqueous layer was dried under N_2_ and resuspended in 1:1 water:acetonitrile. Samples were analyzed by LC/MS using a QExactive benchtop orbitrap mass spectrometer equipped with a heated electrospray ionization (HESI) probe, coupled to a Dionex UltiMate 3000 UPLC system (Thermo Fisher Scientific, San Jose, CA). Samples were separated by injecting 10 μl of each sample onto a ZIC-pHILIC 2.1 x 150 mm (5 μm particle size) column (EMD). Flow rate was set to 100 μL/min, column compartment was set to 25°C, and autosampler sample tray was set to 4°C. Mobile Phase A consisted of 20 mM ammonium carbonate, 0.1% ammonium hydroxide. Mobile Phase B was 100% acetonitrile. The mobile phase gradient (%B) was as follows: 0 min 80%, 5 min 80%, 30 min 20%, 31 min 80%, 42 min 80%. All mobile phase was introduced into the ionization source set with the following parameters: sheath gas = 40, auxiliary gas = 15, sweep gas = 1, spray voltage = ‐3.1kV or +3.0kV, capillary temperature = 275°C, S-lens RF level = 40, probe temperature = 350°C. Metabolites were monitored using a targeted selected ion monitoring (tSIM) method in negative mode with the quadrupole centered on the M-H ion m+1.5, m+2.5, or m+3.5 mass with a 8 amu isolation window, depending on the number of carbons in the target metabolite. Resolution was set to 70,000, full-scan AGC target was set to 106 ions, and tSIM AGC target was set to 105 ions. Relative quantitation of polar metabolites was performed with XCalibur QuanBrowser 2.2 (Thermo Fisher Scientific) using a 5 ppm mass tolerance and referencing an in-house library of chemical standards. Concentration was normalized to cell number.

### Histology

Tissues were fixed overnight to 24 hrs in formalin. Haematoxylin and eosin staining was used for routine histology.

### Quantitation of Melanin in Hair Follicles

HFs with the bulb located entirely in the dermis were considered to be pre-anagen IIIa (specifically anagen II in the case of synchronized samples); HFs with a lower bulb were not included in the analysis. Only HFs with a fully visible bulb were included. Samples were de-identified for blinded quantitation. Each remaining HF was visually assessed for the presence of melanin granules and classified as “none,” “one,” “two” or “three or more.”

### Hair Follicle Synchronization

Mice were anesthetized and plucked over two 1cm^2^ areas halfway down the back, one on each side of the spine, equidistant from the spine. After the procedure, mice were given carprofen at 3 mg/kg once per day for 3 days as an analgesic.

### RT-qPCR

RNA was collected from skin using Trizol reagent (Ambion). Skin samples were digested in 1 mL of Trizol using a GentleMACS tissue homogenizer and RNA was isolated according to standard protocol. RNA from FACS samples was isolated using the RNAqueous Micro Kit (Ambion). cDNA was reverse transcribed using an iScript cDNA Synthesis Kit. RT-qPCR was performed with SYBR Green on a LightCycler 480 II machine from Roche. Primers were used at a final concentration of 1 μM. The primers used are listed in the table below.

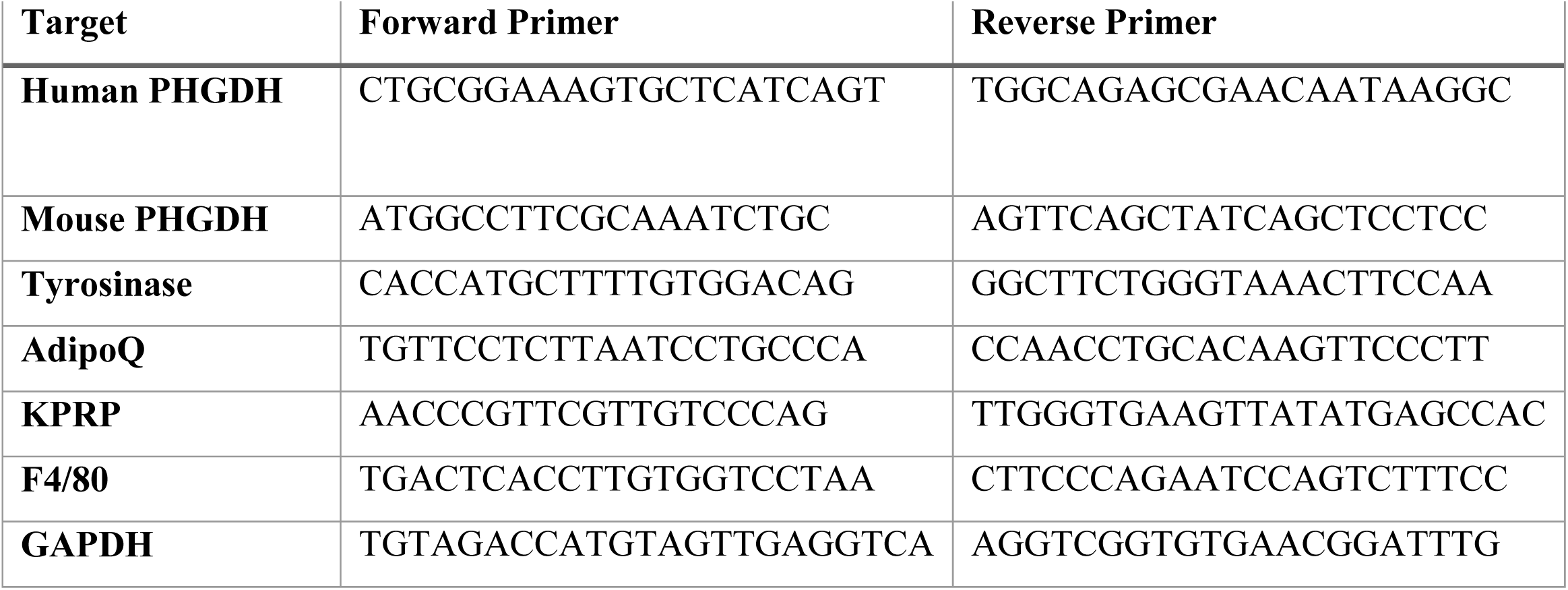

### FACS

Synchronized skin was dissected from mice, then cut into small pieces in a Petri dish using dissecting scissors. The skin was resuspended in 5 mL sterile PBS with 3 mg/mL dispase II (Roche), 1 mg/mL collagenase I (Worthington Biochemical), and 0.1 mg/mL DNase I (Sigma-Aldrich). This solution was incubated at 37 °C for 30 minutes, then EDTA was added to a final concentration of 10 mM to stop the digestion reaction. The digested skin was passed through a 70 μm cell strainer then washed twice with sterile PBS. Cells were stained with 1 μg/mL DAPI for 15 minutes as a live-dead marker, then analyzed for GFP expression on BD FACSAria III flow cytometer.

## ACKNOWLEDGEMENTS

We thank Lauren Surface and Laurie Boyer for their advice on using the flip-in system to generate the *PHGDH*^*tetO*^ mice. We also thank Roberta Ferretti, Jackie Lees, and Marcus Bosenberg for advice on skin sectioning for hair follicles analysis, the Koch Institute Swanson Biotechnology Center for technical support, specifically Aurora Connor and Noranne Enzer at the Mouse ES Cell and Transgenics Core Facility, Kathy Cormier at the Hope Babette Tang (1983) Histology Facility and Scott Malstrom at the Animal Imaging and Preclinical Testing facility. We thank the NCI Mouse Repository for providing us with the iDCT-GFP (01XT4) mice. We acknowledge Michael Pacold for providing recombinant mouse PHGDH enzyme. LC/MS was performed at the Whitehead Institute Metabolite Profiling Core Facility. We thank Jordan Bartlebaugh for performing Western blot analysis. Generation of the *PHGDH*^*tetO*^ mice was supported in part by the MIT Cancer Center support grant (P30-CA14051) from the National Cancer Institute. K.R.M. acknowledges support from the NSF Graduate Research Fellowship Program, DGE-1122374. A.N.L. is a Robert Black Fellow of the Damon Runyon Cancer Research Foundation, DRG-2241-15. K.R.M. and M.R.S. were supported by T32-GM007287. M.R.S. acknowledges support from an MIT Koch Institute Graduate Fellowship. M.G.V.H. acknowledges support from R21-CA198028, R01-CA168653, the Ludwig Center at MIT, SU2C, and a Faculty Scholar Grant from the Howard Hughes Medical Institute.

## SUPPLEMENTAL FIGURE LEGENDS

**Supplemental Figure 1.**
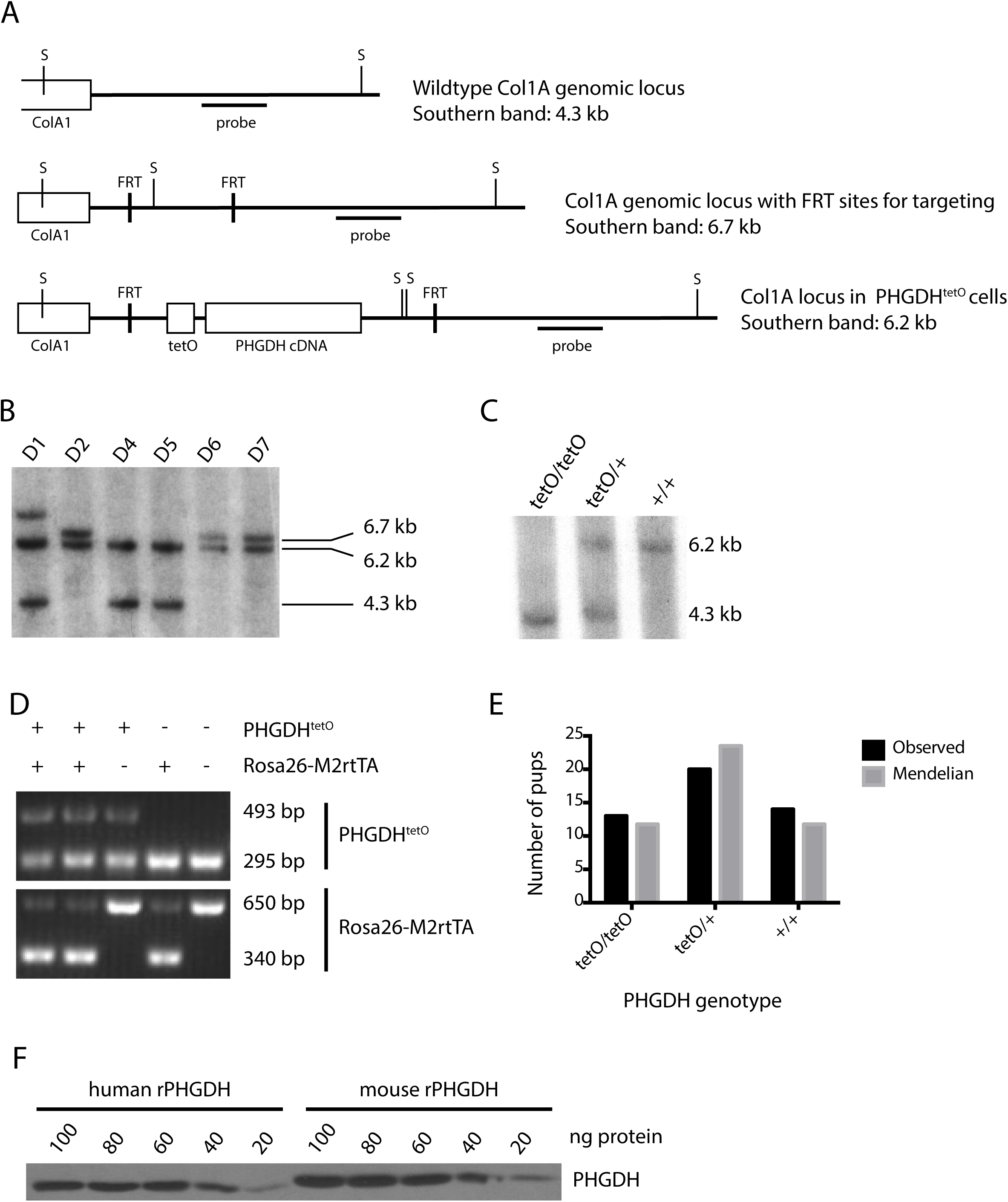
Generation of the *PHGDH*^*tetO*^ allele. **(A)** Schematic of the Col1A locus in wildtype mouse cells (top), the modified locus in KH2 ES cells (middle) and the locus after targeting to introduce the *PHGDH*^*tetO*^ allele (bottom). The PHGDH cDNA introduced into the Col1A locus is the human sequence. The expected band sizes when the indicated probe is used for Southern blot analysis as in **(B)** and **(C)**] are indicated. Also shown are the location of the SpeI sites (marked “S”) in each locus used to digest genomic DNA for Southern blot analysis. FRT, flippase recognition target site; tetO, tetracycline operator minimal promoter. **(B)** Southern blot analysis of SpeI-digested genomic DNA from six *PHGDH*^*tetO*^-targeted ES cells. Clones D4 and D5 exhibit proper targeting of the Col1A locus and an unaffected wild-type allele. **(C)** Southern blot analysis of SpeI-digested genomic DNA from mice of the indicated genotypes. **(D)** PCR-based genotyping of the *PHGDH*^*tetO*^ and *Rosa26-M2rtTA* alleles. In the *PHGDH*^*tetO*^ reaction, the presence of the transgene is indicated by the upper band; in the *Rosa26-M2rtTA* reaction, the presence of the transgene is indicated by the lower band. **(E)** The number of offspring of each genotype observed when mice hemizygous for the *PHGDH*^*tetO*^ allele exposed to a doxycycline diet were mated. The observed distribution of genotypes in the offspring did not differ significantly from expected Mendelian ratios, with p=0.58 by the χ^2^ goodness-of-fit test. **(F)** Western blot analysis of PHGDH protein using the indicated amount of recombinant human or mouse PHGDH to test antibody specificity.

**Supplemental Figure 2.**
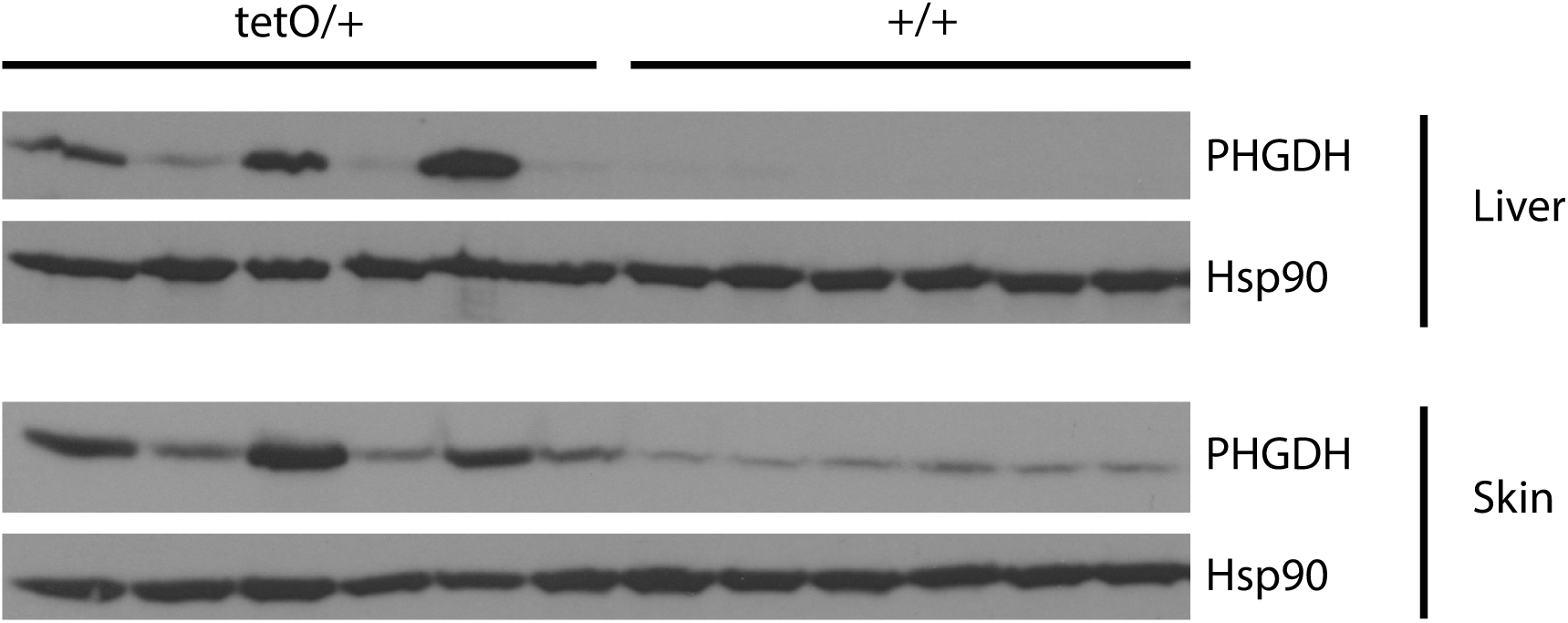
Tissues from mice with long-term exposure to doxycycline diet show variable PHGDH expression. Western blot analysis for PHGDH expression in liver and skin from *PHGDH*^*tetO*^ (tetO/+) and wildtype (+/+) mice that were exposed to doxycycline diet for 16-18 months. Hsp90 expression is also shown as a loading control.

**Supplemental Figure 3.**
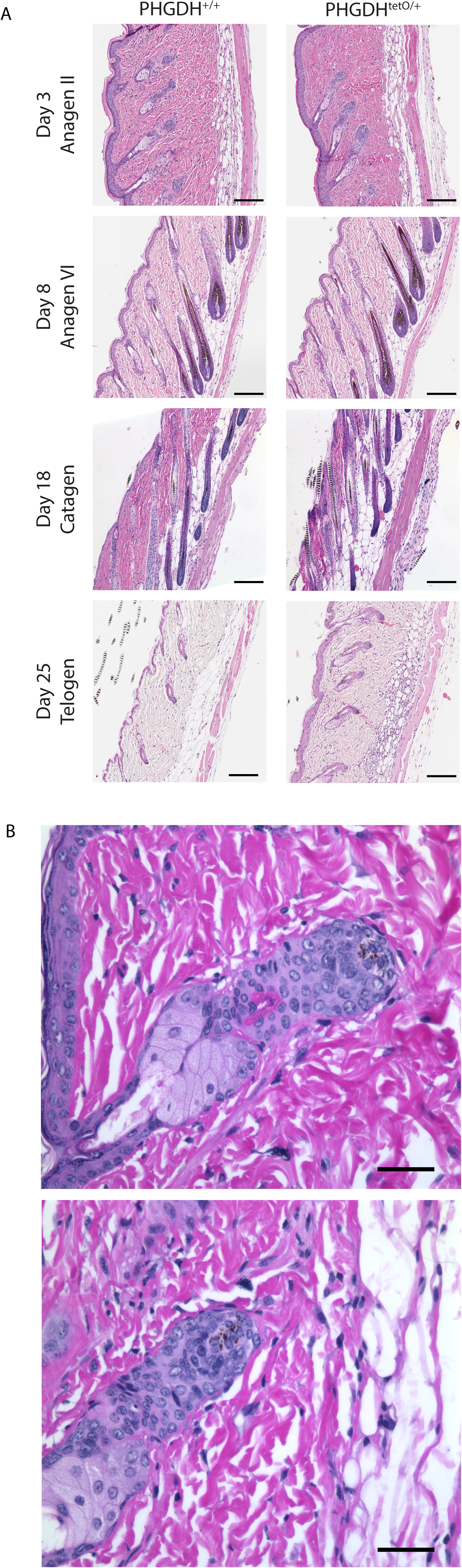
PHGDH expression leads to melanin accumulation in pre-anagen IIIa hair follicles but does not globally affect timing of the hair follicle cycle. **(A)** A region of hair was plucked from 49 day-old mice (at the second telogen) to synchronize the hair follicle cycle, and skin samples were collected at defined days thereafter. Data were collected from *PHGDH*^*tetO*^ or control (+/+) mice that were exposed to doxycycline (Dox) for 30 days prior to synchronization. Representative H&E staining of skin sections from mice of the indicated genotypes is shown. Images were obtained at 4x magnification. Scale bar = 1 mm. **(B)** Representative H&E staining of skin from two *PHGDH*^*tetc*^ *R26-M2rtTA* mice showing anagen II hair follicles (HFs) that contain multiple melanin granules. Images were obtained at 40x magnification. Scale bar = 30 μm.

**Supplemental Figure 4.**
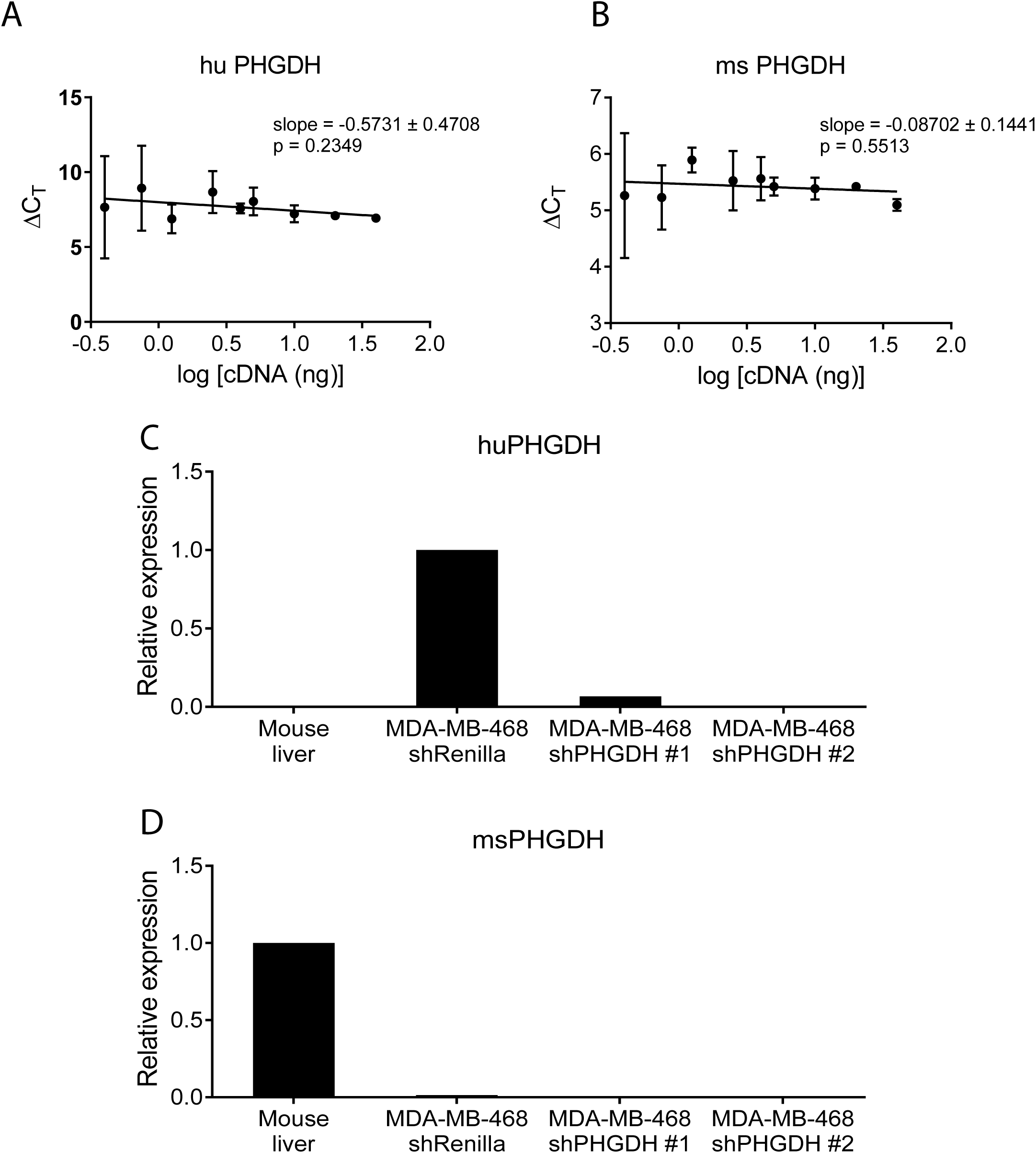
Validation of species-specific PHGDH qPCR primers qPCR primers specific to. **(A)** human (hu PHGDH) and**(B)** mouse (ms PHGDH) PHGDH were tested for linearity and relative quantitation compared to 18S rRNA. The slope of each line does not significantly differ from 0, with p-values derived from an F test. The same human **(C)** and mouse **(D)** PHGDH primers were examined for an ability to amplify PHGDH cDNA derived from mouse liver or the human cell line MDA-MB-468 infected with a control shRNA (shRenilla) or one of two hairpins targeting PHGDH (shPHGDH).

**Supplemental Figure 5.**
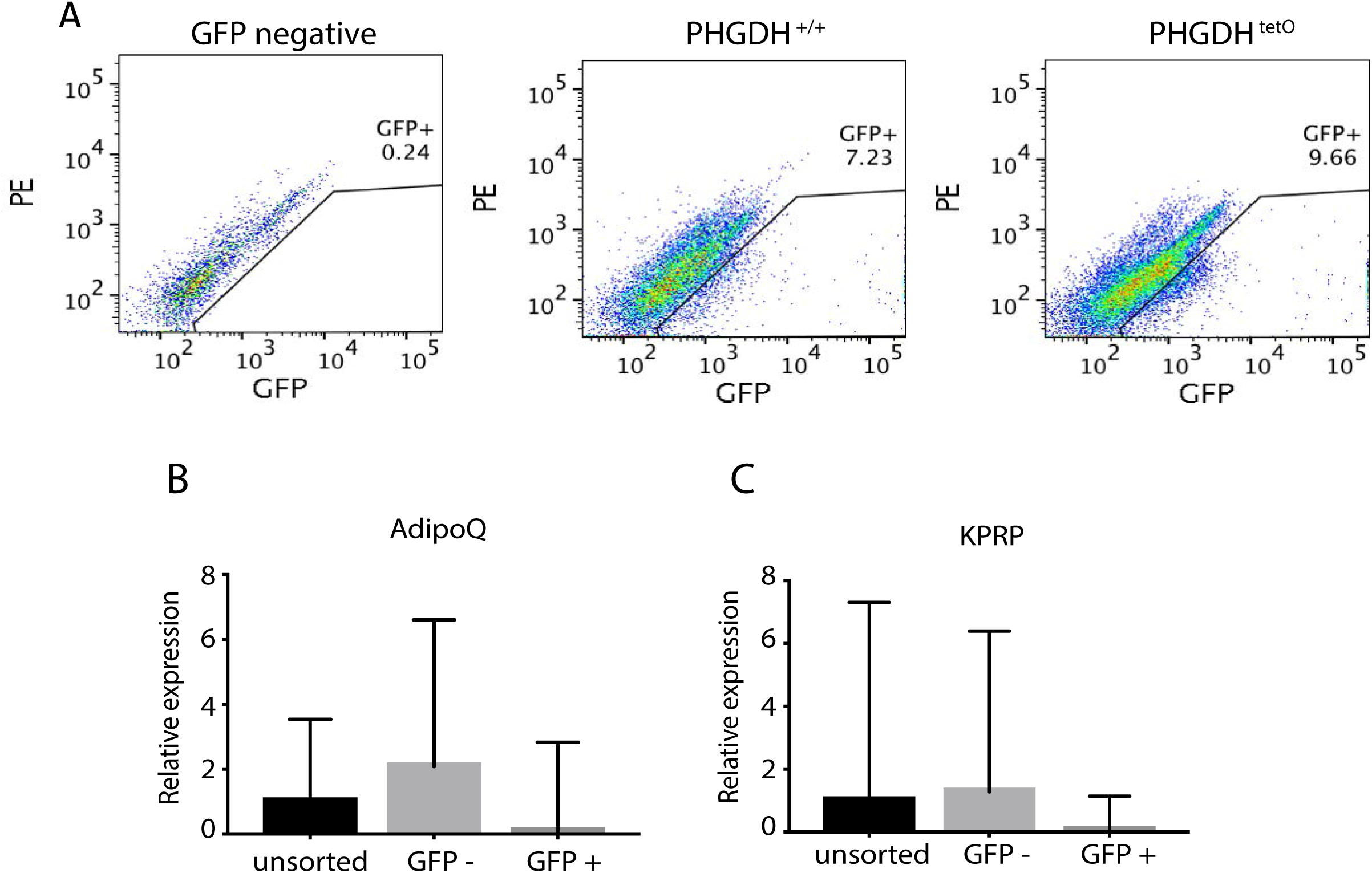
Evidence that adipocytes and keratinocytes sort into the GFP-fraction when cells are isolated from the skin of *PHGDH*^tetO^; *Dct-rtTA*;*H2B-GFP*^*tetO*^ mice. **(A)** *PHGDH*^*tetO*^; *Dct-rtTA*; *H2B-GFP*^*tetO*^ mice with melanocytes that express both PHGDH and GFP, along with control mice with melanocytes that express GFP were exposed to doxycycline for 30 days prior to hair follicle synchronization. Anagen II skin samples were isolated, and GFP+ and GFP-cells were isolated via FACS. Cells were gated on the single cell, live (DAPI-) population. GFP+ cells were identified based on signal in a GFP channel compared to autofluorescence as measured by signal in a PE channel using a skin sample from a mouse that did not contain *H2B-GFP*^*tetO*^ (GFP negative) as a negative control. Representative FACS plots from a PHGDH^tetO^ mouse and a wildtype (PHGDH++) mouse are shown. (B) qPCR analysis of cDNA isolated from unsorted, GFP-, and GFP+ cells isolated from the skin of *PHGDH*^tetO^; *Dct-rtTA*; *H2B-GFP*^*tetO*^ mice described in Figure 4E. Primers were used to amplify the keratinocyte-specific gene *KPRP*, and (C) the adipocyte-specific gene *AdipoQ* as indicated. Data shown represent the mean (+ SD).

